# The Nucleosome Remodelling and Deacetylation complex coordinates the transcriptional response to lineage commitment in pluripotent cells

**DOI:** 10.1101/2023.02.09.527610

**Authors:** Bertille Montibus, Ramy Ragheb, Evangelia Diamanti, Sara-Jane Dunn, Nicola Reynolds, Brian Hendrich

## Abstract

As cells exit the pluripotent state and begin to commit to a specific lineage they must activate genes appropriate for that lineage while silencing genes associated with pluripotency and preventing activation of lineage-inappropriate genes. The Nucleosome Remodelling and Deacetylation (NuRD) complex is essential for pluripotent cells to successfully undergo lineage commitment. NuRD controls nucleosome density at regulatory sequences to facilitate transcriptional responses, and also has been shown to prevent unscheduled transcription (transcriptional noise) in undifferentiated pluripotent cells. How these activities combine to ensure cells engage a gene expression program suitable for successful lineage commitment has not been determined. Here we show that while NuRD is not required to silence all genes, its activity is important to restrict expression of genes primed for activation upon exit from the pluripotent state, and that NuRD activity facilitates their subsequent transcriptional activation. We further show that NuRD coordinates gene expression changes, which acts to maintain a barrier between different stable states. Thus NuRD-mediated chromatin remodelling serves multiple functions, including reducing transcriptional noise, priming genes for activation and coordinating the transcriptional response to facilitate lineage commitment.

## Introduction

Cells in multicellular organisms arise from a single zygote and, with very few exceptions, inherit the same DNA content through cell divisions. Cells in the preimplantation mammalian embryo are totipotent, meaning that they can give rise to all embryonic and extraembryonic cell types of the developing organism. As development progresses cells lose potency as they transit through different states to finally acquire an identity associated with their function. The transitions between the different states are tightly regulated which ensures the correct number of cells is provided for each cell type. This fate acquisition is accompanied by the establishment of an expression profile with specific genes expressed and others actively repressed.

Embryonic stem cells (ES cells) are a powerful system in which to identify differentiation signals involved in cell state transitions. They are highly stable when maintained in the naїve state yet have the potential to form all cell types in an adult organism once allowed to leave that state (1, 2). Indeed, the autocrine signals which drive ES cells to exit pluripotency and initiate lineage commitment, and the cellular machinery important for cells to properly interpret and respond to these signals, have been studied extensively (3–7). Integration of external cues by cells leads to changes in the gene regulatory network which defines cell state, and changes in those cues can further modify the stability of different cell states as development proceeds (8).

Gene expression is controlled to a large extent by how the relevant regulatory sequences - enhancers and promoters - are packaged in chromatin. Chromatin density is largely controlled by protein complexes containing an ATP-dependent catalytic subunit which can move, evict, recruit or assemble nucleosomes (9,10). One such remodeller which is known to facilitate cell fate decisions is the Nucleosome Remodelling and Deacetylase (NuRD) complex (11). NuRD contains both chromatin remodelling and lysine deacetylase activities in distinct subcomplexes (12, 13). These two subcomplexes are held together by the MBD2 or MBD3 proteins, which themselves are mutually exclusive in NuRD complexes (14, 15). In the absence of an MBD2/3 protein the complex falls apart, which does not immediately displace the remodelling component, called CHD4, but it does alter its activity (16). MBD3/NuRD is required for successful lineage commitment of pluripotent cells both in culture and in vivo (17, 18). NuRD components localise to sites of active transcription in multiple cell types, binding to both enhancers and promoters (16, 19–21). Rather than simply activating or repressing transcription, NuRD functions to both dampen unscheduled transcription in the undifferentiated state, and to modulate active transcription as cells are undergoing lineage commitment. Together these transcriptional regulatory activities allow cells to respond appropriately to inductive signals, facilitating lineage commitment (14, 16, 22).

While it is clear that NuRD activity is essential for successful lineage commitment of pluripotent cells, exactly how its transcriptional regulatory activity facilitates lineage commitment is not clear. NuRD activity suppresses inappropriate transcription, or transcriptional noise, in both human and mouse pluripotent cell cultures, but it is not known how this activity is read out in individual cells, nor how cells respond when this noise reduction activity is compromised. Here we used single cell RNAseq across the first 48 hours of ES cell exit from the naїve state to define how Mbd3/NuRD activity functions to facilitate lineage commitment. We determined how NuRD activity impacts the ability of cells to control gene expression as they exit the self-renewing state and identified a function for NuRD in ensuring that cells progress in a coordinated fashion along the early developmental trajectory. This analysis clarifies the role of an important and abundant chromatin remodeller in control of gene expression during lineage commitment, which we show is significantly more nuanced than the now outdated model of remodellers simply switching genes on or off.

## Materials and methods

### Cell lines and differentiation

Mouse embryonic stem (ES) cells were grown on gelatin coated plates in N2B27 supplemented with LIF and the inhibitors PD0325901and CHIR99021 (1) (2iL medium, made in-house). Cell lines were genotyped and tested for mycoplasma contamination regularly. The cell lines used in this study are BHA-derived (WT and *Mbd3*-null; 40, XX) (23) for single cell RNAseq and the bulk of the molecular work and 7E12-derived WT and *Mbd3*-null lines (40, XY) (16) for independent verification of selected results.

Differentiation of embryonic stem cells toward neuroectoderm was induced by withdrawal of two inhibitors and LIF. The cells were grown on laminin coated plates during all differentiation. Differentiation of embryonic stem cells toward mesendoderm was performed on fibronectin coated plates. Cells were grown for 2 days in N2B27 before adding 10 ng/ml activin A and 3 mM CHIR99021 in N2B27 to induce mesendoderm differentiation during subsequent days.

### Single Cell RNA-sequencing

At each time point of the differentiation time course cells were collected using accutase. The resulting cell suspension was sorted using a MoFlo cell sorter (Beckman Coulter) to deposit one cell per well of a 96-well plate in 2 μl of cell lysis buffer (0.2% (V/V) Triton-X100 and 2 U/µL of RNase inhibitor). Library preparation was performed following the Smart-seq2 protocol (Picelli et al., 2014), and libraries were sequenced at the CRUK Cambridge Institute Genomics Core facility (Cambridge, UK) on an Illumina HiSeq4000 sequencer.

### Bioinformatic analyses

RNA-seq libraries were aligned to the mouse reference genome (GRCm38/mm10) with the GSNAP aligner (gmap-2014-12-17) (Wu and Nacu, 2010; Wu et al., 2016) using the parameters -n 1 -N 1 for uniquely mapped reads and allowing for novel splicing sites. Gene read counts were assigned using HTSeq (v0.6.1) based on gene annotation from ensemble release 81. Quality control filtering was then applied to the cells using the R package Scater (1.2.0) (McCarthy et al., 2017) : more that 5*10^5 reads per cell, more than 7000 individual features detected and less than 5% of the reads mapping to mitochondrial genes. R package Seurat v4.1.0 was used for cell filtration, normalization, principal component analysis, variable genes finding, clustering analysis, and Uniform Manifold Approximation and Projection (UMAP) dimensional reduction. The significantly differentially expressed genes were selected at an FDR-adjusted p-value equal or lower than 0.05. The gene ontology analysis was conducted using MouseMine (Motenko et al., 2015).

### Discretization of the data and fate score analysis

The discretization of the expression data for individual genes was conducted using k-mean clustering into two groups except for the genes with unimodal distribution of expression. According to the clustering the cells were classified as low expressing cells or high expressing cells for each gene. The fate score for each cell was calculated using representative genes for every lineage and summing the number of genes with high expression for these genes. Representative pluripotency genes were *Pou5f1, Sall4, Zfp42, Klf2, Sox2, Tfcp2l1, Stat3, Nanog, Klf4*, and *Tbx3* (24, 25). Representative neuroectoderm genes were *Wnt8a, Cxcl12, Enpp2, Bcl11a, Sema6a, Pbx1, Pax6*, and *Sox1*. The correlation analysis was conducted using Pearson correlation test. Cell-cell correlation was conducted by computing the Pearson coefficient of correlation between each pair of cells based on the gene expression of all the genes detected. This was done individually at each time point per genotype.

### ChIP-qPCR

ChIP was performed exactly as described (26). ChIP was performed with anti-FLAG antibody (Sigma, F1804) using 10 µg/ChIP, and with Rabbit IgG at the same concentration. Locus-specific primers used for quantitative PCR are listed in the Supplementary Table.

### MNase qPCR analysis

Cells were harvested at the mentioned time point using accutase and washed once in ice cold PBS. One million cells were next resuspended into 1mL of ice-cold lysis buffer (10 mM Tris-HCl pH7.4, 10 mM NaCl, 3 mM MgCl2, 0.5% NP-40 (Igepal), 150 mM spermine and 500 mM spermidine). The suspension was centrifuged at 300 × g for 10 min at 4°C and the pellet washed in 1 ml digestion buffer (10mM Tris-HCl pH7.4, 15mM NaCl, 60mM KCl, 150mM spermine and 500mM spermidine). MNase digestion was performed in 100 µl of digestion buffer containing 1 mM CaCl and 40 U of Mnase (New England Biolabs) at 24°C for 15min while shaking. The reaction was stopped by adding 100 µL of stop buffer (digestion buffer containing 20 mM EDTA, 2 mM EGTA). Samples were next treated with RNase and Proteinase K. Monosomal DNA was extracted using phenol chloroform and gel purification after gel electrophoresis. Locus-specific quantitative PCR was used to quantify the signal across the region of interest. The signal was normalised using sonicated genomic DNA and the average signal of the entire region. Locus-specific primers used for quantitative PCR are listed in the Supplementary Table.

### Gene Expression Analysis

Total RNA was purified using Trizol reagent following the manufacturer instructions. DNase treatment was performed using DNase I and First-strand cDNA was synthesised using SuperScript IV reverse transcriptase (Invitrogen) with random hexamers. Quantitative PCRs (qPCRs) were performed using TaqMan® reagents (Applied Biosystems) and SybrGreen® (Applied Biosystems) on a QuantStudio Flex Real-Time PCR System (Applied Biosystems). Gene expression was determined relative to housekeeping genes (*Gapdh, Atp5A1, Ppia*) using the ΔCt method. Locus-specific primers used for quantitative PCR are listed in the Supplementary Table.

## Results

### MBD3/NuRD activity is required for mouse ES cells to express lineage-appropriate genes

To define the average behaviour of Mbd3/NuRD mutant ES cells during the early stages of lineage commitment, we investigated gene expression changes by RT-qPCR which occur as cells leave the naїve state and begin differentiating down the neuroectodermal lineage. In wild type cells pluripotency-associated genes (*Tbx3, Klf4, Nanog* and *Zfp42*) were rapidly downregulated, while neuroectoderm markers (*Pou3f1, Sox1, Irx3* and *Pax6*) were activated steadily across a 72 hour differentiation time course (Figure 1A). By contrast, while *Mbd3* mutant cells downregulated the pluripotency-associated genes, they failed to induce the expression of lineage genes by 72 hours (Fig 1B). This same result was seen in two independently derived *Mbd3*-null ES cell lines (Supp Fig 1). To determine if the defect in the induction of the expression of lineage genes was limited to neuroectoderm genes, we conducted a similar analysis during mesendoderm differentiation (Fig 1C). Here again, the *Mbd3* mutant cells failed to properly induce the expression of lineage genes after 4 days (*T, Sox17, Mixl1* and *Eomes*) whereas the downregulation of the pluripotency genes was not affected by the absence of *Mbd3* (Fig 1D). These results are consistent with our previous results differentiating ES cells or primed human and mouse pluripotent cells lacking NuRD activity (14, 22), and indicate that a fully functional NuRD complex is required for cells to respond properly to the differentiation signals and induce expression of lineage-specific genes.

**Figure 1.**
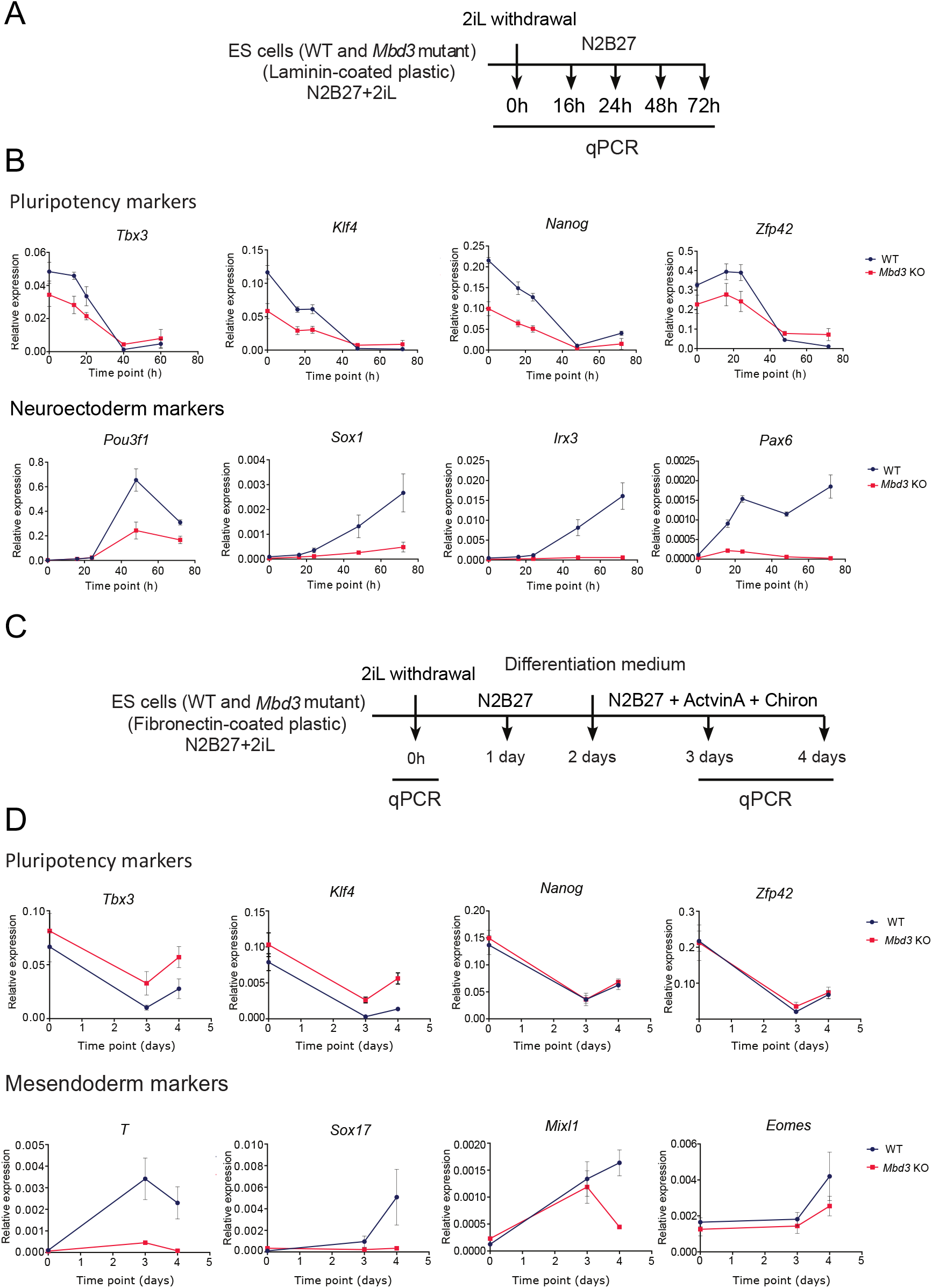
Gene expression analysis at the population level confirms the defect of induction of lineage genes during differentiation in *Mbd3* mutant cells. **A**. Protocol for monolayer neuroectodermal differentiation of naїve ES cells in N2B27 (25). **B**. Gene expression analysis by RT-qPCR at the population level for selected representative pluripotency markers and neuroectoderm markers during neuroectoderm differentiation. Expression was normalised to three housekeeping genes (*Gapdh, Atp5A1, Ppia*). Error bars indicate the standard error of 4 independent differentiations. **C**. Protocol for mesendoderm differentiation of naїve ES cells in N2B27 supplemented with Activin A and Chiron (27). **D**. Gene expression analysis by RT-qPCR at the population level for selected representative pluripotency markers and mesendoderm markers during mesendoderm differentiation. Error bars indicate the standard error of 4 biological independent differentiations.

### NuRD binding and nucleosome remodelling activity at regulatory regions of lineage-specific genes correlates with their induction

NuRD regulates gene expression changes during early differentiation through chromatin remodelling activity at promoters and enhancers (Bornelöv et al., 2018). To verify that induction of lineage-specific gene expression during differentiation is associated with NuRD-dependent chromatin changes, we first asked whether MBD3/NuRD is associated with regulatory regions of differentiation-associated genes. We evaluated MBD3 binding at enhancer and promoter regions of two genes normally induced during neuroectodermal differentiation of ES cells (*Irx3* and *Pax6*) by ChIP-qPCR (Supp Fig 2A). Enrichment of MBD3 increases during differentiation both at enhancer and promoter regions of the two lineage genes tested (Fig 2A and Supp Fig 2B). This is consistent with our observation that NuRD associates with transcriptionally active sites (16). This increase in MBD3 enrichment coincided with an increase in nucleosome positioning at discrete sites within two different enhancers as measured by MNase protection (position 16672 for *Pax6* and -80441 for *Irx3*; Fig 2B, D). In the absence of MBD3 this increased positioning does not occur, consistent with this being a NuRD-dependent remodelling event. By contrast, the other nucleosomes in the Pax6 enhancer either become more positioned (nucleosome at ∼15800) or shift to a new position (nucleosome at ∼14500) in mutant cells upon differentiation. Notably, chromatin remodelling activity was focussed at enhancer regions as, though the corresponding promoter regions showed a differentiation-associated increase in MBD3 enrichment, we detected no significant changes in nucleosome density at promoter regions (Supp Fig 2B). Together these data show that increased MBD3 enrichment and MBD3-dependent nucleosome remodelling at distal regulatory regions of lineage-specific genes coincides with differentiation-induced gene activation.

**Figure 2.**
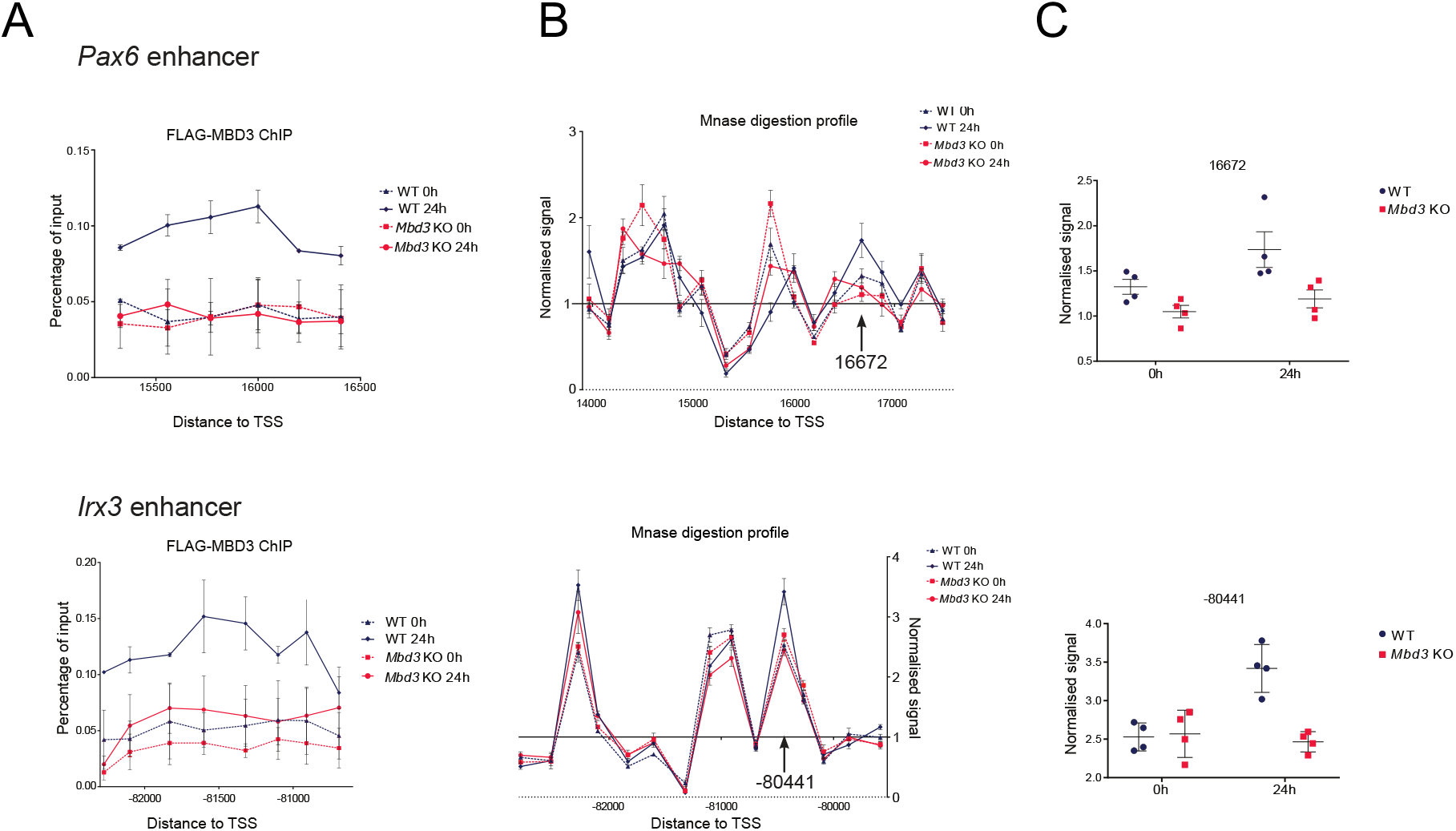
Binding of MBD3 and NuRD dependent chromatin remodelling activity at distal regulatory regions confirm the direct role of NuRD in the induction of lineage-specific gene expression. **A**. ChIP-qPCR for MBD3 (FLAG) across the *Pax6* and *Irx3* enhancers before differentiation (dashed lines) and after 24 hours of differentiation (plain lines) in WT (blue) and *Mbd3* mutant cells (red). x axis show locations relative to the annotated transcription start site (TSS). Error bars indicate the standard error of 3 (0h) and 2 (24h) independent differentiations. **B**. Mnase-qPCR data across the *Pax6* and *Irx3* enhancer before differentiation (dashed lines) and after 24 hours of differentiation (solid lines) in WT (blue) and *Mbd3* mutant cells (red). The x-axes show locations relative to the annotated transcription start site. The arrow indicates the position where changes were observed after 24 hours only in WT cells and further analysed in C. Error bars indicate the standard error of 4 independent differentiations. **C**. Detail of the normalised signal at the positions 16672 and -80441 for the enhancer of *Pax6* and *Irx3* respectively. Individual experiments are shown, and the horizontal bars indicate the mean and the standard error of 4 independent differentiations. While the difference in MNase protection between WT and Mbd3 KO is not significant at time 0, for both genes the difference is significant at 24 hours (p < 0.03, Mann-Whitney test).

### NuRD facilitates a robust and uniform transcriptional response to differentiation cues

NuRD activity is required to suppress transcriptional noise as pluripotent cells initiate lineage commitment, and cells lacking NuRD activity can respond to differentiation cues but cannot establish the correct gene regulatory networks required for successful lineage commitment (8, 14, 22). To explore the role of NuRD in control of gene expression during lineage commitment in more detail, we assayed mRNA expression in individual wild type or *Mbd3*-null ES cells in self-renewing conditions (2iL) and at 12, 24 and 48 hours after removal of both inhibitors and LIF (Fig 3A). We chose to use SMRT-seq rather than a more high-throughput method as we wished to maximise the depth of mRNA recovery to better detect lowly expressed genes. We therefore sequenced 96 cells per time point and per genotype, making a total of 768 cells, 739 of which passed quality control and were used for subsequent analyses (Supp Fig 3A). To validate the approach and the single cell sequencing we looked first at the expression of genes identified as early responders after 2iL withdrawal (Kalkan et al., 2017; Sokol, 2011). As expected, we observed an increase in the expression of ERK-responsive genes and a decrease in the expression of WNT target genes during differentiation of the WT cells (Supp Fig 3B).

**Figure 3.**
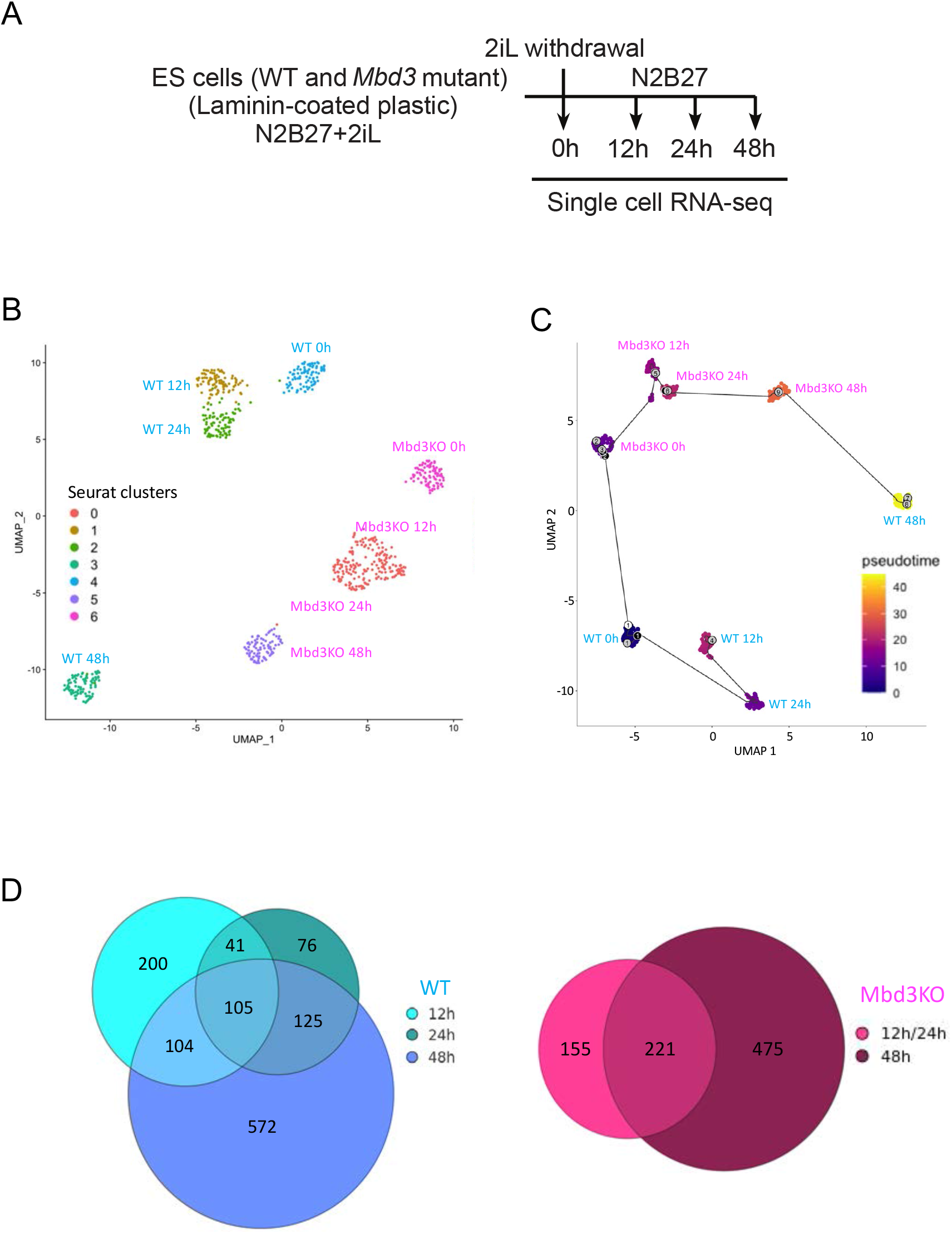
Single cell gene expression analysis of exit from the naїve state in wild type and *Mbd3*-null ES cells. **A**. Scheme of differentiation time course for the single cell RNA sequencing dataset. **B**. UMAP representation of the single cell RNAseq data using all genes. Only cells that have passed quality control are plotted. Cells are coloured according to their Seurat cluster affiliation and labelled with their biological identities. **C**. Pseudotime analysis showing trajectories of cells according to their distance from WT 0h cells and coloured according to their distance to the WT 0h state. **D**. The numbers of significantly (P < 0.05) differentially expressed genes in each Seurat clusters (reassigned to their corresponding biological condition) are shown for wild type (left) and *Mbd3*-null cells (right).

Visualising the data as a UMAP representation shows that wild type and mutant cells cluster separately throughout the differentiation time course (Fig 3B). Despite clustering separately at the 2iL time point, both wild type and mutant cells show movement through the UMAP plot across the differentiation time course in the same general direction. This indicates that NuRD activity is not required for cells to receive and begin to respond to differentiation cues. Yet the mutant cells only show limited movement across the UMAP plot (Fig 3B). Further, while WT cells were separated into 4 clusters corresponding to differentiation time using the Seraut algorithm (see Methods), the same method could not bioinformatically distinguish between the 12 and 24 hour time points for mutant cells (Fig 3B, D). Pseudotime analysis (28) similarly showed that mutant cells are arranged along a differentiation trajectory, but do not proceed as far as wild type cells after 48 hours (Fig 3C).

Comparison of the number of differentially expressed genes (DEGs) at every time point revealed that most of the changes occur by 48 hours of differentiation in both genotypes (Fig 3D). Enrichment analysis for gene ontologies on DEGs at 48 hours indicated mostly general developmental terms and terms associated with metabolism and motility, with neural development being the predominant developmental lineage as expected when 2iL is removed from ES cell culture (29) (Supplementary Table 1). Genes activated in mutant cells at 48 hours were also associated with general developmental, metabolism and motility terms, although less significantly than wild type cells (Supplementary Table 2).

The GO term “Nervous system development” was enriched in both the WT and in the mutant after 48 hours of differentiation (WT: P= 9.56E-06; KO P = 2.85E-02). We were surprised at this, given that *Mbd3*-null cells have an extremely low probability of successfully adopting a neuroectodermal fate, even in the correct embryonic context (18). We therefore compared the behaviour of genes within this GO term between the wild type and mutant differentiation time courses (Fig 4). As a class, these genes are expressed in wild type cells at very low levels in 2iL and remain low through 24 hours. Between 24 and 48 hours of differentiation these genes go from low expression to being much more decisively on or off. In *Mbd3*-null cells these genes are similarly expressed at low levels for the first 24 hours of differentiation, and while some genes do show increases or decreases in expression at 48 hours, the change is far less pronounced or robust as in wild type cells. This is consistent with NuRD activity not being necessary for cells to either produce or recognise a signal inducing strong activation or silencing of genes associated with “Nervous system development,” but rather NuRD activity is required for cells to maintain the appropriate transcriptional response to these signals.

**Figure 4.**
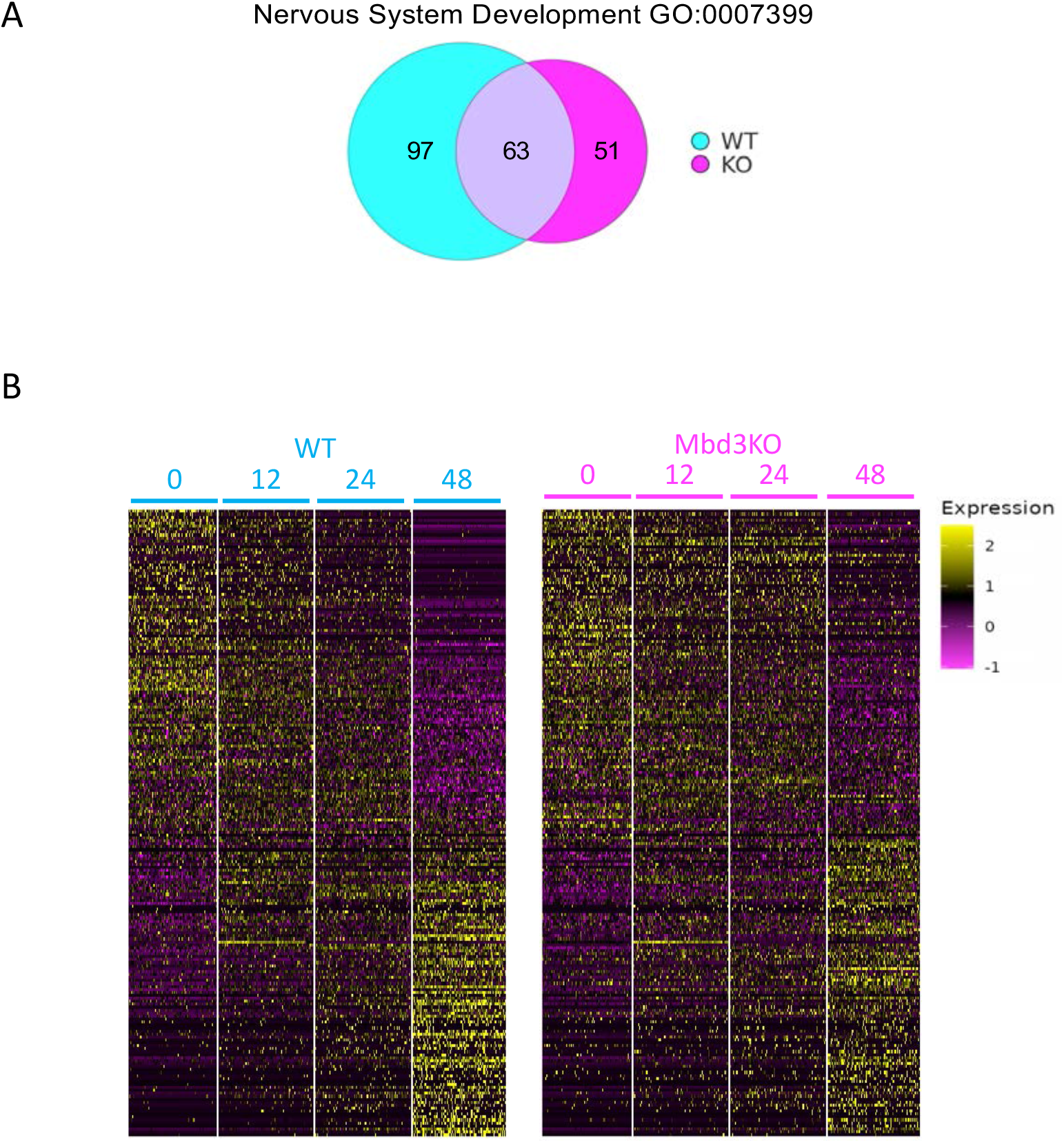
Expression of differentially expressed neural genes across the differentiation time course. **A**. Venn diagram showing the overlap between genes enriched under the term “Nervous system development” (GO:0007399) for WT (blue) and Mbd3-Null cells (magenta). **B**. Heatmap showing the log2 normalised expression level for all differentially expressed genes under the GO term “Nervous system development”.

### Mbd3/NuRD maintains coordination of differentiation-induced gene expression changes

To get a picture of the behaviour of changes in the neural and pluripotency gene regulatory networks across the differentiation time course, we calculated a “fate score” per cell by classifying cells using k-means clustering in two groups according to their expression (high or low) for a set of representative pluripotency-associated genes (see Materials and Methods). A score of 1 was given to cells with high expression and 0 for cells with low expression, and these used to calculate a global score per cell for groups of genes. These “fate scores” at different time points in WT and mutant cells mimicked the results obtained with normalised gene expression levels (Supp Fig 4A). Distribution of pluripotency scores among cells showed that, by 48 hours, there were 9 cells with the maximum pluripotency score (10) in the mutant cells, compared to none in the WT cells (Fig 5A). Moreover, we observed higher heterogeneity in the distribution of the pluripotency score for the mutant cells after 48 hours as compared to the WT cells which switch quickly from a high to low score between 24 and 48 hours (Fig 5A). This indicates that NuRD activity normally maintains the integrity of the gene regulatory network during differentiation, such that in the absence of Mbd3/NuRD cells variably deviate from the normal developmental trajectory leaving some able to initiate early developmental gene expression, some unable to leave the pluripotent state, and the rest at some point between these two extremes.

**Figure 5.**
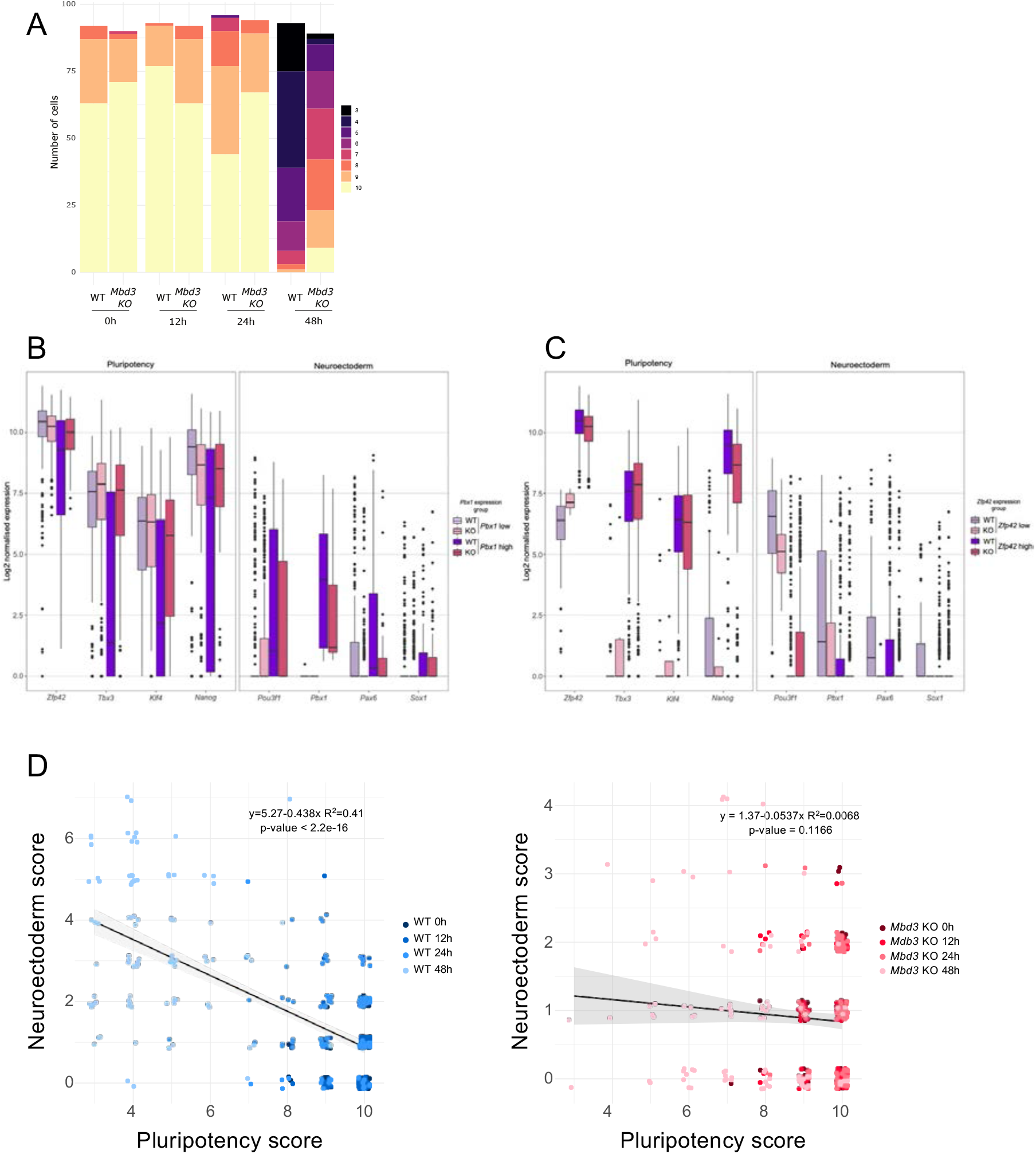
NuRD is important for a coordinated transcriptional response to exit from the naїve state. **A**. The cumulative number of cells for each pluripotency score is plotted for wild type and *Mbd3*-null cells at each time point. Colours indicate pluripotency scores. **B**. The distribution of the normalised expression of indicated genes, representative of pluripotency (left) or neuroectoderm (right) after grouping the cells according to their expression level (low in purple or high in pink) of *Pbx1*. **C**. Same as panel B but cells grouped for expression of *Zfp42*. **D**. Pearson correlation analysis between the pluripotency and neuroectoderm fate scores for the WT (blue) and *Mbd3* mutant single cells (red). The grey area represents the 5% confidence interval of the linear regression.

The decrease in expression of pluripotency genes but lack of, or reduced activation of, differentiation-associated markers in *Mbd3*-null ES cells after 48 hours in differentiation conditions indicates that there is a lack of coordination of differentiation-associated gene expression changes in the absence of NuRD activity. To investigate this in more detail we firstly discretised expression levels for an early neural marker, *Pbx1* (30), as well as for a marker of naїve pluripotency, *Zfp42*. We classified cells as *Pbx1*-high or low across all time points and plotted cells according to expression of selected pluripotency and neuroectoderm genes (Fig 5B, C). In WT cells expression of *Pbx1* is associated with low levels of pluripotency gene expression and activation of early differentiation genes, as expected for differentiating cells. By contrast, in the mutant cells this coordination of gene expression changes is partially lost, with *Pbx1* high or low cells showing little difference in the expression of pluripotency markers, even though *Pbx1* high cells show some activation of other neuroectodermal markers (Fig 5B). If we instead classify cells in terms of expression of *Zfp42*, a marker of naїve pluripotency, *Mbd3*-mutant *Zfp42*-low cells show both lower expression of neural markers and elevated expression of naїve markers *Tbx3* and *Klf4* as compared to wild type *Zfp42*-low cells (Fig 5C). These data indicate a lack of coordination between silencing of the pluripotency gene expression network and activation of the neural differentiation programme in the absence of Mbd3/NuRD.

To obtain a broader view of the coordination of gene expression changes between the pluripotency and the neuroectoderm networks, we used the calculated fate scores to conduct a correlation analysis (Fig 5D). Whereas the WT cells showed the expected negative correlation between the pluripotency score and the neuroectoderm score (Pearson correlation, r = -0.63674, p<2.2e-16), *Mbd3* mutant cells showed a lack of any significant correlation (Pearson correlation, r = -0.08228127, p= 0.1166) (Fig 5D). NuRD activity is therefore required for coordination of the transcriptional response induced upon exit from the self-renewing state.

### Incomplete silencing of differentiation-associated genes precedes failure of their activation

To define how NuRD-dependent gene regulation defects change over time, we identified those genes significantly changing expression in wild type cells at each time point during the differentiation time course and tracked how they behave in mutant cells. Firstly, it was notable that all gene sets analysed showed a broader range of expression in self-renewing conditions in wild type cells than in mutant cells (Fig 6B). This indicates that NuRD activity allows cells to maintain a greater dynamic range of expression of these gene subsets in naїve ES cells. Secondly, Mbd3/NuRD was not generally required for gene downregulation at 12 or 24 hours, but at 48 hours mutant cells showed persistent expression of genes which were effectively switched off in wild type cells (e.g. 24 and 48 hour DOWN genes, Fig 6B). Thirdly, while genes normally activated at all three time points show increases in expression in *Mbd3*-null cells, in all three cases the level of activation is reduced compared to that seen in wild type cells. For example, those genes normally activated at 12 hours show both an increased average expression and more homogeneous expression in undifferentiated mutant cells. After 12 hours in differentiation conditions these genes show an increase in average expression levels, but do not reach the levels seen in wild type cells, and subsequently do not get effectively downregulated by 48 hours (Fig 6A, B). Together these data indicate that NuRD activity increases the transcriptional dynamic range of naїve pluripotent ES cells, and is important both for transcriptional activation as cells exit the naїve state and for complete repression of genes normally silenced as cells transit through formative pluripotency.

**Figure 6.**
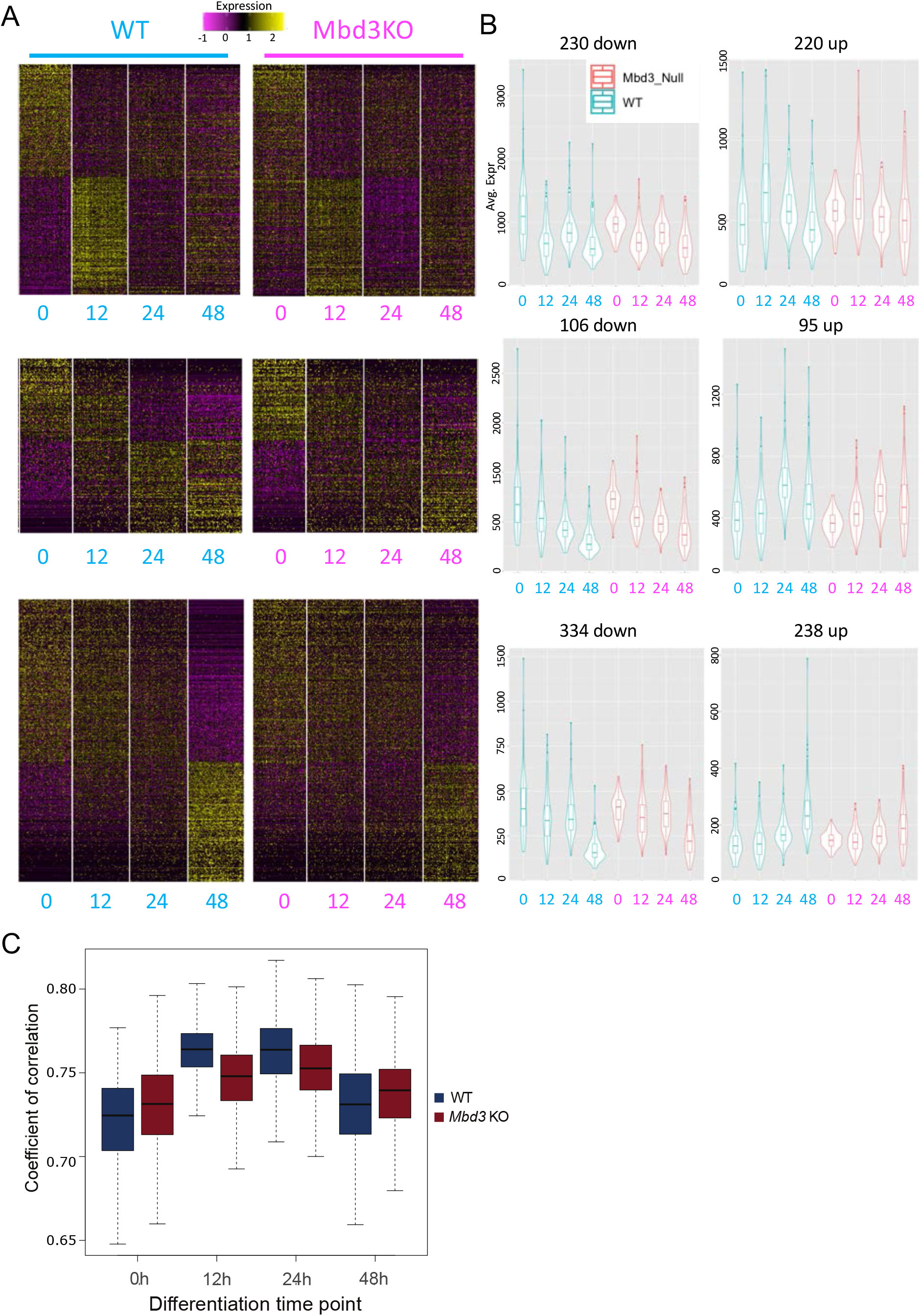
NuRD activity is required both for noise reduction and for timely activation of specific gene subsets. **A**. Average expression of genes significantly changing in wild type cells plotted across all time points, clustered according to fold change in wild type cells. Expression of genes first showing significant changes at 12 hours (top panels), 24 hours (middle panels) or 48 hours (bottom panels) are displayed for wild type (left) and *Mbd3*-null cells (right). **B**. Average expression profiles of gene sets shown in A, with wild type expression patterns shown in blue (left) and expression in mutant cells shown in red (right). The shape of the violin indicates the proportion of cells at each position along the y-axis; the midline of the box shows the median value, the lower and upper edges of the box represent the 1st and the 3rd quartiles, and the whiskers represent the minimum and maximum values. **C**. Distribution of the Pearson coefficient of correlation between the WT (blue) and *Mbd3* mutant cells (red) is shown at each time point of differentiation.

Mbd3/NuRD has been shown to suppress transcriptional noise in both human and mouse pluripotent cells (14, 22). To explore this finding further we calculated the pairwise correlation between cells of a given genotype and time point. Indeed *Mbd3*-null cells showed an increased cell-cell correlation (P-value < 2.2e-16; Welch’s t-test) in self-renewing conditions compared to WT cells (Fig 6C). The initial response to withdrawal of 2iL is relatively uniform, causing a large increase in the cell-cell correlation in both wild type and mutant cells. In mutant cells however, the transcriptional response, and the increase in cell-cell correlation, is reduced compared to that seen in wild type cells. By 48 hours correlation between wild type cells is reduced as cells begin to adopt different fates, while the defective lineage commitment in *Mbd3*-null cells leaves them more homogeneous.

We therefore conclude that the essential function exerted by NuRD to facilitate lineage commitment is two-fold: firstly, it reduces transcriptional noise globally in undifferentiated conditions, and secondly, it is required to mount an appropriate, coordinated transcriptional response to receipt of differentiation signals.

## Discussion

Here we provide clarification of the essential role played by the NuRD complex in control of gene expression during ES cell lineage commitment. We find that NuRD is important both for keeping the early responding differentiation-associated genes repressed in the self-renewing state, and for their timely and efficient activation upon 2iL withdrawal. We suggest that this is because NuRD activity is required to render the regulatory sequences of these genes appropriately responsive to activation signals. It is notable that NuRD does not appear to be required for active downregulation of highly transcribed genes in ES cell differentiation, which must be achieved by some other repressive mechanism. Rather, our data support a model in which NuRD activity is necessary both to prevent precocious gene activation and to effectively prime genes for scheduled activation.

The ability to profile gene expression changes in individual cells allowed us to obtain a detailed picture of cellular behaviour during differentiation. While it is clear that NuRD-mutant cells are capable of downregulating components of the pluripotency gene regulatory network (Fig 1) (14, 22), after 48 hours of differentiation we can still identify some *Mbd3*-null cells with persistent expression of pluripotency markers (Fig 5A). This explains why it was possible to recover self-renewing *Mbd3*-null ES cell colonies several days after differentiation induction (Reynolds et al. 2012). This indicates that the probability of completely inactivating the pluripotency gene regulatory network is reduced in the absence of NuRD activity. It also indicates that cells read this probability at least somewhat independently such that in a culture of millions of cells, the vast majority will have silenced the pluripotency gene regulatory network, but a small proportion have not. By contrast NuRD activity in wild type cells ensures that the probability of remaining pluripotent after exit from the self-renewing state is effectively zero. This difference, between a low probability and a near-zero probability, is important because persistence of even one pluripotent cell in a developing somatic lineage would result in teratoma formation.

NuRD proteins localise to regulatory regions of active genes in multiple cell types and CHD4-dependent nucleosome remodelling has been identified at both enhancers and promoters (16, 19–22, 31, 32). Yet at the genes investigated in detail here, we detected NuRD-dependent nucleosome movements at enhancers, but not promoters, after 24 hours in differentiation conditions (Fig 2, Supp Fig 2b). Enhancers are likely to be the important drivers of changes to gene expression during development (7), so control of chromatin structure by NuRD at enhancers is likely to directly control their ability to respond to inductive signals. How chromatin remodellers and extracellular signals combine to alter enhancer chromatin has not been systematically investigated.

In addition to controlling access of transcription factors and the general transcription machinery to regulatory sequences, NuRD also functions to increase the space explored by enhancers in the nucleus (16, 33). Whether these two activities are directly related remains to be determined. Nonetheless, we propose a scenario where enhancers of genes primed to respond to signals are held in an inactive but activatable state through NuRD-mediated chromatin remodelling, which restricts access of transcription factors to enhancer chromatin (Fig 7). Upon receipt of a differentiation signal these enhancers, which are in the correct chromatin configuration to rapidly become activated, can interact with relevant promoters and instruct gene expression changes. In the absence of NuRD, differentiation-associated enhancers lack the appropriate protein binding repertoire to fully function. Upon receipt of a differentiation signal, some responsive gene promoters become activated though the relevant enhancers, while others are unable to contact appropriate enhancers and hence are not activated appropriately. In the absence of a coordinated, robust transcriptional response lineage commitment is not possible.

**Figure 7.**
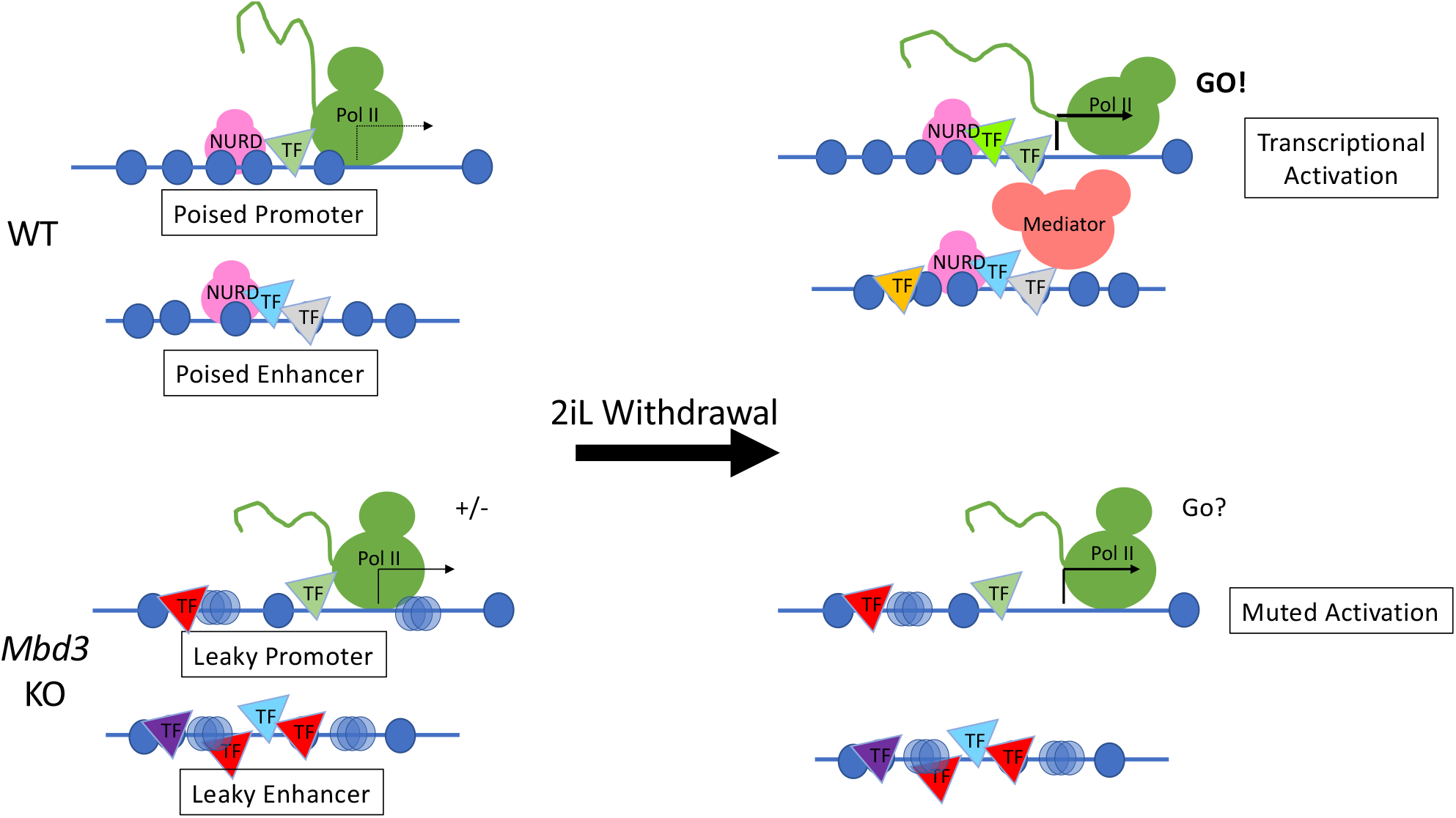
Model of how NuRD facilitated transcriptional activation during differentiation. In wild type cells NuRD activity ensures positioned nucleosomes (blue circles) and restricts transcription factor (triangles) binding at promoters and enhancers (top left). The promoter is poised, but inactive. The enhancers are very mobile in 3D space, meaning they can scan a large area for sequences with which to interact. When cells are induced to differentiate (2iL Withdrawal), NuRD maintains enhancer nucleosome structure and allows appropriate transcription factor binding so the enhancer and promoter can interact to drive differentiation-associated transcription (top right). In *Mbd3*-null cells both promoters and enhancers have less positioned nucleosomes (hazy blue circles), inappropriate transcription factor binding (bottom left), and the enhancer is less mobile, meaning it can only sample its immediate vicinity. The lack of NuRD activity at the enhancer and/or promoter results in incomplete silencing (“+/-”). When induced to differentiate the enhancer is not in an appropriate chromatin configuration an cannot sample sufficient 3D space to appropriately activate transcription. As a consequence the promoter shows muted activation (bottom right).

## Data Availability

The single cell RNA-seq datasets reported in this study are available from the Gene Expression Omnibus (GEO) repository under accession code GSE214264 (wild type cells) and XXXXX (*Mbd3* KO cells).

## Acknowledgments

We thank Céline Labouesse and members of the Hendrich, Laue, Arnaud and Oakey labs for discussion, advice, and comments on the manuscript. We thank Andy Riddell, Maike Paramour and Sally Lees for expert technical assistance and advice.

## Author Contributions

BM, NR SJD and BH designed the research, BM generated the data, BM and RR performed bioinformatics analyses, BM, RR and BH made the figures, BM and BH wrote the manuscript with input from other authors and BH, NR and SJD obtained funding.

---

**Supplementary Figure 1.**
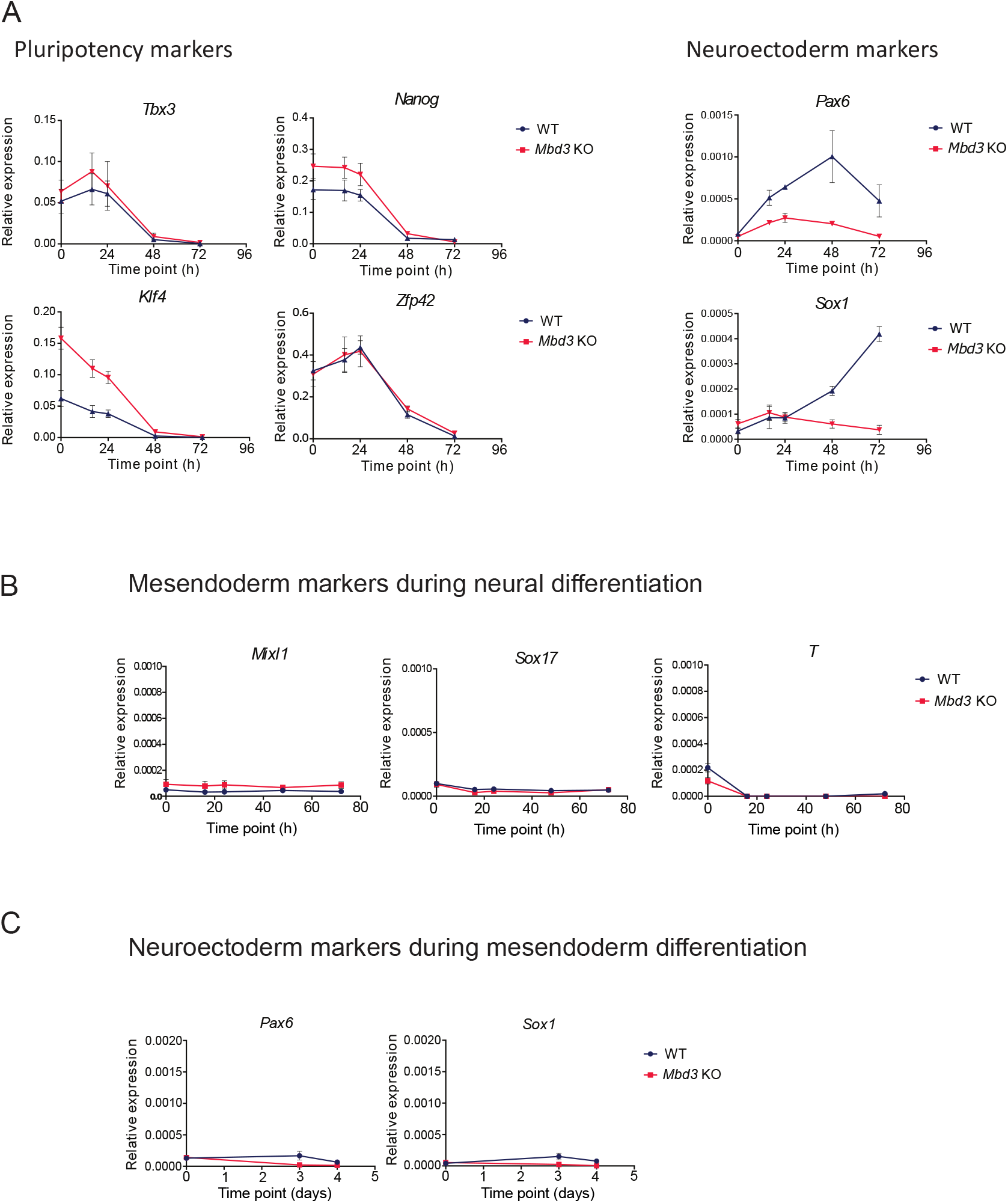
*Mbd3* mutant cells show a defect of induction of lineage gene expression and an absence of other lineages activation during neuroectoderm differentiation. **A**. Gene expression analysis by RT-qPCR at the population level for selected representative pluripotency markers and neuroectoderm markers during neuroectoderm differentiation of another *Mbd3* mutant cell line. Error bars indicate the standard error of 4 independent differentiations. **B**. Gene expression analysis by RT-qPCR at the population level for selected representative mesendoderm markers during neuroectoderm differentiation. Error bars indicate the standard error of 4 independent differentiations. **C**. Gene expression analysis by RT-qPCR at the population level for selected representative neuroectoderm markers during mesendoderm differentiation. Error bars indicate the standard error of 4 independent differentiations.

**Supplementary Figure 2.**
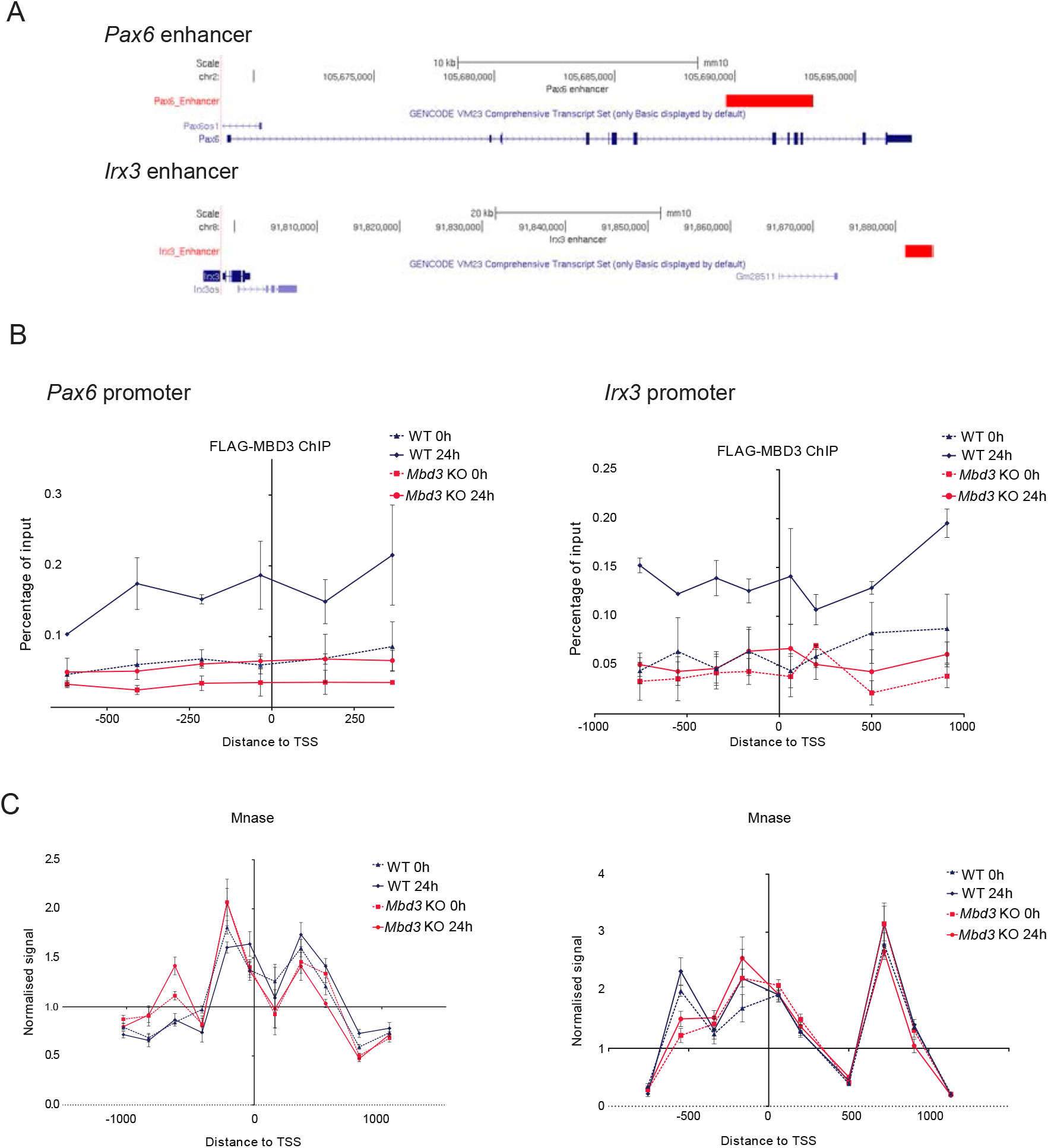
Recruitment of MBD3 at promoter regions is not associated with differentiation specific chromatin remodelling activity at promoter regions. **A**. UCSC genome browser capture (mm10 assembly) of the *Pax6* and *Irx3* loci studied in this paper. The red boxes represent the enhancer regions analysed. **B**. ChIP-qPCR for MBD3 (FLAG) across the *Pax6* and *Irx3* promoters before differentiation (dashed lines) and after 24 hours of differentiation (plain lines) in WT (blue) and *Mbd3* mutant cells (red). The x axes show locations relative to the annotated transcription start site. Error bars indicate the standard error of 3 (0h) and 2 (24h) independent differentiations. **C**. Mnase-qPCR data across the *Pax6* and *Irx3* promoters before differentiation (dashed lines) and after 24 hours of differentiation (plain lines) in WT (blue) and *Mbd3* mutant cells (red). The x axes show locations relative to the annotated transcription start site. The arrow indicates the position where changes were observed after 24 hours only in WT cells and further analysed in C. Error bars indicate the standard error of 4 independent differentiations.

**Supplementary Figure 3.**
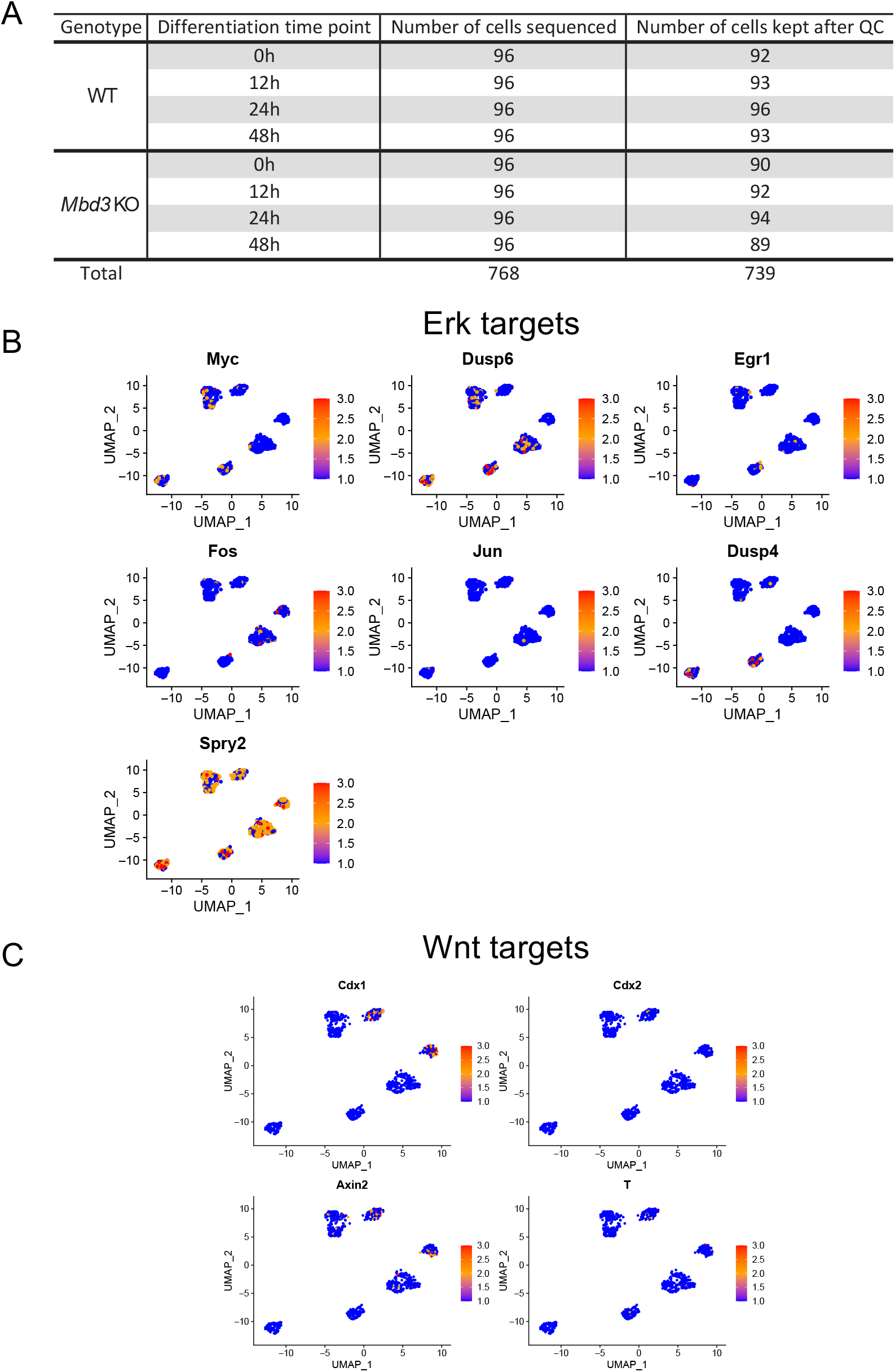
Differential expression analysis of the single cell RNA-seq dataset supports the proper differentiation of the WT cells and the upregulation of the placenta development associated genes in the *Mbd3* mutant cells. **A**. Table summarising the number of cells sequenced and the cells which passed quality control. **B**. Heatmap showing the log2 of the normalised expression level of genes identified as ERK response genes and WNT target genes which are early responders of the 2iL withdrawal (25), in the WT cells. **C**. Venn diagram showing the number of genes differentially expressed in WT (blue) and *Mbd3* mutant cells (red) at 12, 24 and 48 hours of differentiation in comparison to 0h.

